# Microbiota induces aging-related leaky gut and inflammation by dampening mucin barriers and butyrate-FFAR2/3 signaling

**DOI:** 10.1101/2021.08.18.456856

**Authors:** Sidharth P Mishra, Bo Wang, Shaohua Wang, Ravinder Nagpal, Brandi Miller, Shalini Jain, Jea Young Lee, Cesar Borlongan, Subhash Taraphdar, Sushil G. Rane, Hariom Yadav

**Affiliations:** USF Center for Microbiome Research, University of South Florida Morsani College of Medicine, Tampa, FL, USA; Department of Neurosurgery and Brain Repair, University of South Florida Morsani College of Medicine, Tampa, FL, USA; Center for Excellence of Aging and Brain Repair, University of South Florida Morsani College of Medicine, Tampa, FL, USA; Department of Nutrition and Integrative Physiology, Florida State University, Tallahassee, FL, USA; Department of Chemistry, North Carolina A&T State University, Greensboro, NC, USA; Department of Animal Genetics and Breeding, West Bengal University of Animal & Fishery Sciences, Kolkata, West Bengal, India; Diabetes, Endocrinology and Obesity Branch, National Institute of Digestive and Diabetes and Kidney Diseases, National Institutes of Health, Bethesda, MD, USA; Department of Internal Medicine- Digestive Diseases and Nutrition, University of South Florida Morsani College of Medicine, Tampa, FL, USA

**Keywords:** Microbiome, aging, gut, barrier, mucus, permeability, inflammation

## Abstract

Increased chronic inflammation is one of the key risk factors of aging-related disorders although its precise etiology remains elusive. Here, we demonstrate that aged, but not young, microbiota triggers inflammation by promoting gut permeability (leaky gut) via disruption of mucus barriers. Levels of the beneficial short-chain fatty acid, butyrate, are suppressed in the aged gut. Consistent with feedback regulation, the expression of butyrate-sensing receptors, free fatty acid receptor 2/3 (FFAR2/3), are also reduced in aged gut. Butyrate treatment of aged mice revereses the reduced mucin production, increased gut permeability and inflammation associated with low butyrate levels. In agreement, intestine-specific FFAR2/3 knockout mice manifest a compromised gut phenotype typically seen in aged mice,, such as increased gut permeability and inflammation with reduced mucin production. Taken together, our results demonstrate that an aged gut microbiota causally instigates inflammation by increasing gut permeability due to reduced butyrate levels, FFAR2/3 expression, and mucin barriers. Thus, butyrate-FFAR2/3 agonism could ameliorate the deleterious effects seen in aged gut and their implications on metabolic health.

## INTRODUCTION

With the increasing aging and veteran population, the prevalence of aging-related diseases, such as diabetes, obesity, cardiovascular dysfunctions, Alzheimer’s disease, and cancer, is on the rise (Disease et al., 2018). Geroscience is an emerging field focused on understanding the basic biology of aging, which may lead to improve the clinical interventions, as well as included inflammation as one of the pillars of Geroscience and may lead to the generation of clinically meaningful outcomes to improve health of aged adults (Ferrucci & Fabbri, 2018; Kennedy et al., 2014). Persistent low-grade chronic inflammation is often observed in older adults and is considered one of the major risk factors of aging-related diseases (Ferrucci & Fabbri, 2018). The persistent decline in adaptive immunity and chronic increase in the levels of cytokines like interleukin (IL)-1β, IL-6, tumor necrosis factor-alpha (TNF-α) in the systemic circulation are hallmarks of increased inflammation in older adults (Koelman et al., 2019). However, the precise origins of chronic inflammation in elderly are obscure. We and others have shown that gut permeability (referred to as ‘leaky gut’) is often increased in older adults and is an understudied source of inflammation in aging (Ahmadi, Wang, et al., 2020; Wilms et al., 2020). Increased gut permeability allows the non-specific diffusion of microbes, microbial ingredients, metabolites, and antigens from the gut lumen to the blood which, in turn, can promote the migration of pro-inflammatory molecules and systemic inflammation. For example, lipopolysaccharide (LPS) -one of the cell wall components of gram-negative bacteria - leaks from gut causing systemic inflammation resulting in endotoxemia (Cani et al., 2008). Indeed, levels LPS-binding protein (LBP) and soluble CD14 (sCD14) – markers of leaky gut - are significantly higher in the older adults and are linked with age-associated phenotypes such as decreased physical activity and increased risk of heart-failure (Awoyemi et al., 2018; Stehle et al., 2012). However, the mechanisms underlying compromised gut barrier functions in the aged gut are unknown.

Gut barrier functions are under the purview of two major structural components: mucin and tight-junction proteins (Chelakkot et al., 2018). Mucin forms a physical barrier by sealing gaps between the intestinal epithelial cells, while tight junction proteins tightly adhere epithelial cells. Intestinal goblet cells produce mucin as part of a thick gel like viscous mucus layer on the intestinal epithelia, thus facilitating the selective diffusion of small molecules from the gut lumen to intestinal epithelial cells (Pelaseyed et al., 2014). Although this mucus layer restricts any direct bacterial interactions with intestinal epithelial cells, it allows the colonization of bacteria on the mucuosal surface (Li et al., 2015). Mucin also serves as a food source for numerous gut bacteria such as the mucin-degrading bacteria *Akkermansia* (Liu et al., 2021). Mucin depletion, however, permits easy interaction of microbes with intestinal epithelial cells, thus initiating degradation of tight junctions proteins, increasing gut permeability and inflammation in the gut environment (Okumura & Takeda, 2018). Mucin 2 (Muc2) and mucin 6 (Muc6) are the major mucin isoforms in the intestine (McGuckin et al., 2011). In agreement, Muc2-deficient mice develop severe colitis and gut inflammation, thereby highlighting the physiological importance of mucin in maintaining gut barrier functions and gut permeability while staving off inflammation (Van der Sluis et al., 2006). Indeed, aging rodent and primate models exhibit thin mucin lining linking increased gut inflammation with age-associated diseases (Elderman et al., 2017; Sovran et al., 2019; Vemuri et al., 2020). Taken together, maintaining a healthy mucus layer is critical for ideal gut barrier function with advancing age. However, the mechanisms and molecular drivers of reduced mucin levels in the aging gut are poorly understood.

The gut microbiota in older adults is distinct compared to young adults (Langille et al., 2014). Specifcally, aged gut microbiota signature is often characterized by skewing of the ratio in favor of increased abundance of harmful bacteria and reduced proportions of beneficial bacteria. Termed dysbiosis these perturbations to the gut microflora dysregulate the production of adverse and beneficial metabolites (Ahmadi, Wang, et al., 2020). However, evidence that fluctuations in the microbiota population instigate gut permeability and promote gut inflammation in elderly adults is lacking. It is also unknown if the gut microbiota impact goblet cells and mucin barriers in the gut of elderly adults. Furthermore, the mechanisms by which microbiota impact the mucus and tight junction barrier functions are unclear. Using un-biased, non-supervised and comprehensive analyses, we address these open questions. We unravel causal microbiota-dependent mechanisms with translational potential to reverse gut permeability and inflammation in aged adults and help mitigate the consequential adverse health outcomes.

## RESULTS

### 1. Aged gut microbiota propogates gut permeability and inflammation

Microbiome diversity is an important indicator of microbiota balance and health in the gut (Lozupone et al., 2012). The β-diversity indicates similarity between the two microbiome communities (a higher value indicates more differences between the microbiomes), while the α-diversity indicates the microbial diversity within the samples (a higher α-diversity indicates more diversity) (Wagner et al., 2018). Principal co-ordinate analysis (PCA) of β-diversity showed that the gut of aged mice displayed significantly (p<0.05) distinct microbiota signatures than that from younger mice (Figure 1a). The α-diversity indices like Shannon index and number of operational taxonomic units (OTUs) were lower in the aged gut compared to the younger gut (Figure 1b, Supplementary Figure S1a), suggesting that the aged microbiome skewed towards unhealthy state. Furthermore, compared to the younger gut the abundance of major phyla such as Bacteroidetes, Verrucomicrobia, Proteobacteria, Deferribateres and Actinobacteria were lower in the aged gut while the Firmicutes abundance was high (Figure 1c, Supplementary Figure S1b). Similar differences in the old and young gut were observed in other taxonomic ranks, including class, orders and family (summarized in Supplementary Figure S1c-e). Furthermore, the abundances of major genera like *Barnesiella, Dysgonomas, Akkermansia, Lactobacillus, Parabacteroides, Alistipes, Turicibacter* and *Bacteroides* were significantly lower in the aged gut, while the abundances of *Clostridium_XIVa, Acetobacteroides* and *Lachnospiracea* genera were higher (Figure 1d). Similar results were obtained at the genera level by both heatmap and linear discriminatory analysis (LDA) effect size (LEfSe) analysis (Supplementary Figure S1f,g). At the species level, we also found that the abundance of *Muribaculum intestinale, Akkermansia muciniphila, Turicibacter sanguinus, Acetatifactor muris, Mucispirillum schaedleri, Lactobacillus intestinalis, Barnesialla intestinihominis* and *Faecalibaculum rodentium* species were significantly lower, while *Bacteroidetes bacterium, Eubacterium cellulosolvens* and *Flintibacter butyricus* were higher in old gut compared to young (Figure 1e). LEfSe analyses also revealed similar results at species level (Supplementary Figure S1h) with the aged gut harboring a significantly distinct microbiome signature on different taxonomic levels like unique phyla, class, order, family and genera (Figure 1f). In addition, we also find that the aged mice have significantly higher (p<0.01) gut permeability (measured by increased transfusion of FITC [Fluorescein isothiocyanate]-dextran from the intestine to systemic blood circulation) and increased mRNA expression of inflammatory markers (IL-1β, IL-6 and TNF-α mRNA expression) (Figure 1g,h). Moreover, the concentration of systemic inflammatory markers like IL-6 and TNF-α concentrations were significantly higher (p<0.01) in the aged mice as compared to the younger mice (Figure 1i,j). These results indicate that aged gut harbors a microbiota distinct from that seen in the younger gut, with the former linked with increased gut permeability and systemic inflammation. We then inquired whether the microbiota differences are causal in the acquisition of aging-related increases in gut permeability and inflammation.

**Figure 1.**
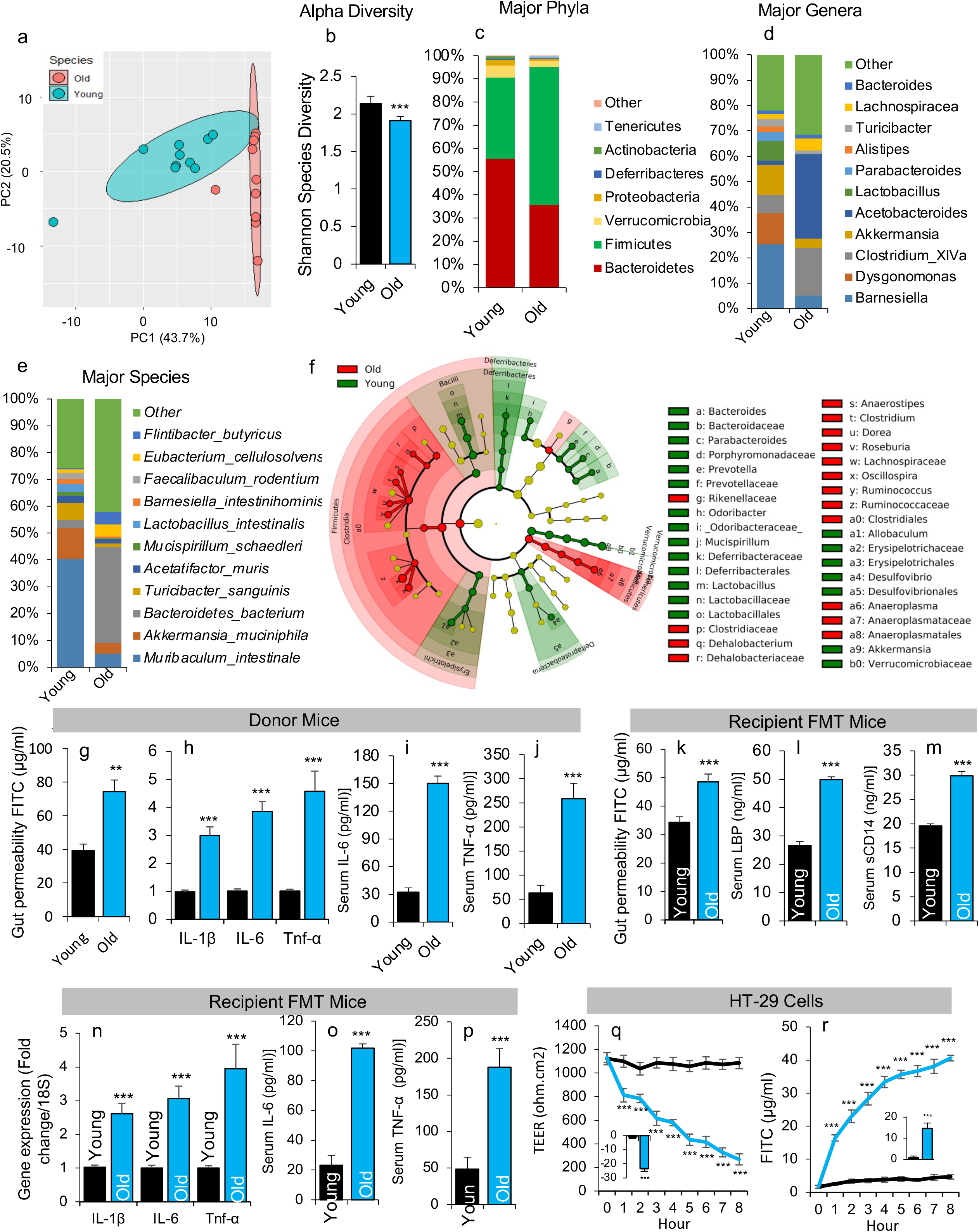
An aged gut harbors significantly distinct microbiota than a younger gut, which is causal to induce leaky gut and inflammation. a) Principal component analysis (PCA) as a β-diversity indices revealed that aged gut microbiome are significantly distinct from a younger gut. b) The microbiome diversity α-diversity indices indicated by the Shannon index was significantly lower in feces from an aged gut as compared to a young gut. c-f) The abundance of major phyla, genera and species presented in bar-graphs (c-d) and Linear discriminant analysis (LDA) effect size LEfSe cladogram is significantly distinct in an aged microbiome compared to a younger microbiome. g-j) Aged mice tend to have an increased amount of leakiness from the gut as represented by FITC-dextran leakage from gut to blood, and increased expression of IL-1β, IL-6 and TNF-α) in the gut along with higher circulating IL-6 and TNF-α as compared to a younger gut. k-p) FMT of aged feces significantly increased leaky gut (FITC-dextran leakage) and its systemic markers like LBP and sCD14 in serum along with increase in inflammatory markers (IL-1β, IL-6 and TNF-α) in gut and IL-6 and TNF-α in serum of recipient mice compared to young FMT recipients. q-r) Treatment of FCM made from aged gut feces significantly reduced transepithelial electrical resistance (TEER) and increased FITC-dextran permeability in the monolayers of HT-29 cells compared to FCM of younger gut feces. All the values presented are mean of 5-9 animals in each group and error bars represent standard error of means. P-values with **p<0.01, ***p<0.001 are statistically significant.

Fecal microbiota transplantation (FMT) from aged mice significantly increased FITC-dextran diffusion and elevated levels of systemic leaky gut markers like LBP and sCD14 in young recipient mice (Figure 1k-m), suggesting that aged microbiota triggers leaky gut in recipient young mice. In addition, recipient mice with aged FMTs also showed higher levels of systemic inflammation markers like Il-1β, Il-6 and Tnf-α mRNA expression in the intestine and the serum (IL-6 and TNF-α) compared to the young FMT controls (Figure 1n-p). These findings suggest that the aged microbiota can initiate gut permeability and inflammation. Furthermore, we showed that, compared to FCM of younger mice, the treatment of human intestinal epithelial HT-29 cell monolayers with fecal conditioned media (FCM) of aged mice significantly decreased the trans-epithelial electrical resistance (TEER) and increased the diffusion of FITC-dextran (Figure. 1q,r). These observations also suggest that the aged microbiota instigates leakiness in the gut epithelia, which in turn stimulates inflammation. Further, these results demonstrate that FCM treatments recapitulate the effects of FMTs using a cell culture system and, thereby, are alternatives to FMTs to study the impact of the microbiota on epithelial barrier function. Overall, these results indicate that the gut of aged mice harbors an abnormal microbiome with reduced microbial diversity, which elevates gut permeability and inflammation.

### 2. Aged microbiota induces gut permeability and inflammation by disrupting mucus barriers

FMT of aged mice significantly reduced mRNA expression of mucin (Mucin 2 [Muc2] and mucin [Muc6]), tight junction genes (tight junction protein 1 [Tjp1] and occludin [Ocln]) expression in the gut of older FMT recipient mice compared to their young FMT recipient controls (Figure 2a,b). Similarly, expression of gut barrier markers was reduced in intestinal organoids and intestinal goblet cells (CMT93 cells) upon incubation with aged mice FCM (Figure 2c-e). To understand the mechanism of action, we next performed unsupervised correlation analyses (heat-map), random forest analyses (RFA) and differential expression (Volcano graph) via gene expression and ELISA from mice, organoids and CMT-93 cells. Muc2 expression exhibited the highest negative correlation with markers of leaky gut, inflammation and comprised intestinal barrier (Figure 2f-h). This suggests that mucin may be the primary target of aged microbiota to induce gut permeability. Histological analyses also showed that the number of goblet cells were significantly decreased in mice receiving aged mice FMT, compared to those receiving young mice FMT (Figure 2i-j). These data indicate that the aged microbiota reduced the abundance of goblet cells, accompanied by lower Muc2 expression and poor intestinal barrier functions. Further, the gut of aged donor mice displayed significantly less expression of Muc2, Muc6, Tjp1 and Ocln, along with reduced goblet cell numbers and lower mucin levels in their feces compared to young mice (Figure 2k-o). Thus, the aged mice manifest compromised gut barrier functions due to a reduced number of goblet cells, which is linked to decreased mucin production. Taken together, these results indicate that aged microbiota promotes gut permeability by disrupting intestinal barrier functions via reduction in goblet cell numbers and decrease in mucin production.

**Figure 2.**
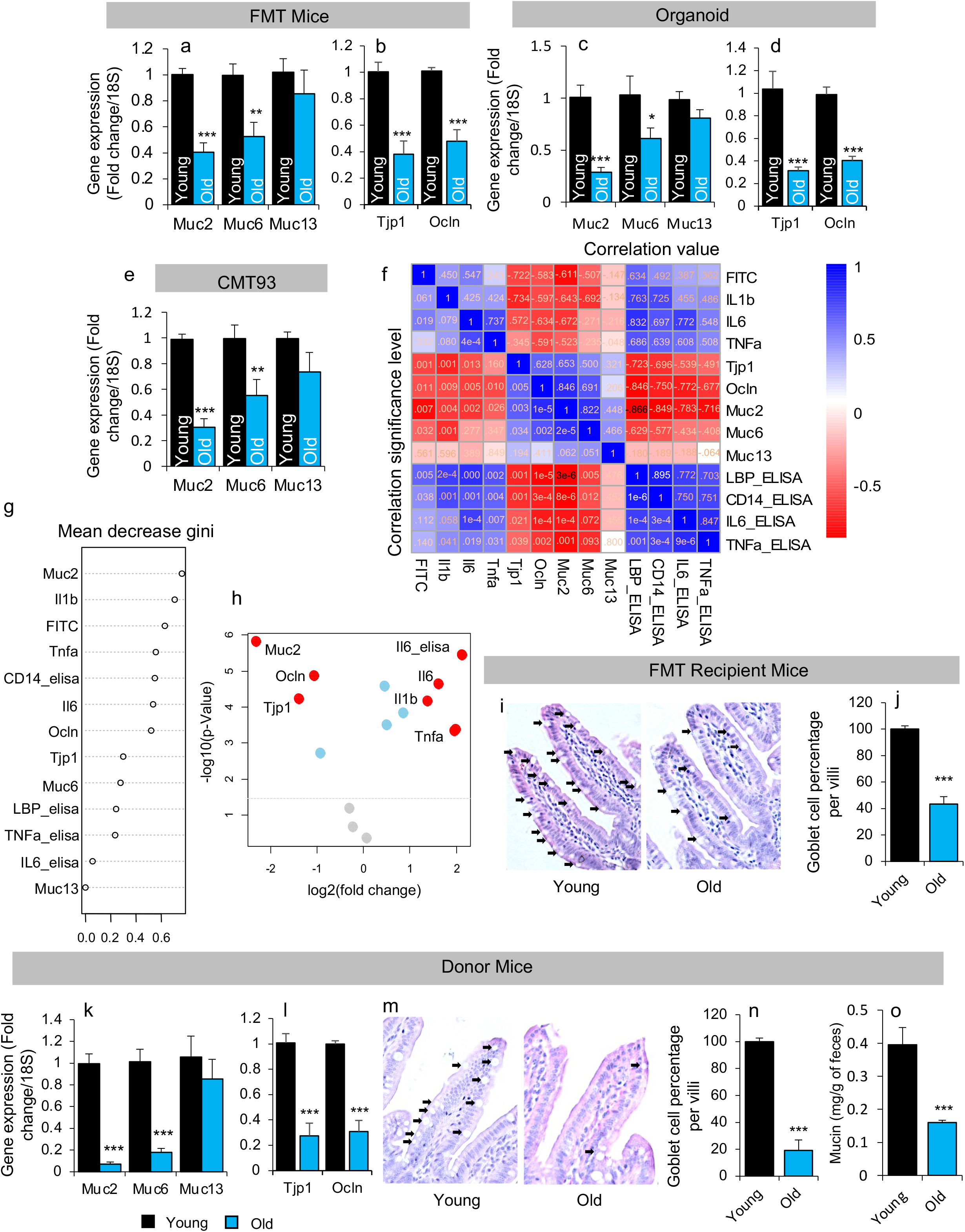
Aged gut microbiota disrupts gut mucin barriers, which in turn induces leaky gut and inflammation. a-e) The expression of mucin isomers like Muc2 and Muc6 (a,c,e), and tight junction protein like Tjp1/zonulin and Ocln (b,d) genes was significantly reduced in the gut of old FMT recipient mice (a,b) and old FCM treated organoids (c,d) and CMT93 goblet cell line (e) in comparison to their corresponding controls. f-h) Pearson’s correlation heatmap (f), random forest analysis (g) and volcano plot (h) representing data of mucin, inflammation and gut permeability markers in the gut of aged FMT recipient mice and FCM treated organoids and CMT93 cells shows Muc2 was the highest influenced gene from an old microbiome. i-j) Similarly, goblet cell numbers were significantly lower in the intestine of old FMT recipients than young FMT recipient controls. k-o) the expression of mucin (k), tight junction proteins (l), goblet cell numbers (m,n) and mucin in feces (o) were significantly lower in aged mice as compared to younger mice. All the values presented are mean of 5-9 animals in each group and error bars represent standard error of means. P-values with *p<0.05, **p<0.01, ***p<0.001 are statistically significant.

### 3. Reduced short-chain fatty acids production in aged gut is linked to reduced mucosal barrier and increased gut permeability and inflammation

Muc2 expression shows the highest negative correlation with gut permeability and inflammatory markers (Figure 3a). This further suggests that reduced mucin production is a primary manifestation in aged donor mice, instigated by an abnormal microbiota. Considering that metabolites produced by gut microbiota are key mediators that impact host cells (Hariom Yadav, 2016), we next performed metabolomics analysis to determine the mechanism by which gut microbiota impacts Muc2 expression. PCA revealed that the feces of aged mice had significantly different metabolic signatures compared to the feces of young mice (Figure 3b). Further, differential analyses showed that the abundance of beneficial metabolites such as short-chain fatty acids (SCFAs like acetate, propionate and butyrate) and taurine were significantly reduced in the aged gut while abundance of anserine, total bile acids, cholesterol and cholate were significantly increased (Figure 3c). Un-biased RFAs showed that the SCFAs (like butyrate and propionate) were most significantly reduced in the aged gut compared to the younger gut (Figure 3d). Among the metabolites, levels of butyrate were the most significantly reduced in the the aged gut (Figure 3e, and Supplementary Figure 2a). Correlation analyses also showed that the reduced butyrate levels were linked to the reduced Muc2 expression and markers of increased gut permeability and inflammation (Figure 3f-h, Supplementary Figure 2b). Together, these findings are consistent with the notion that the aged microbiota reduces butyrate production, thereby decreasing mucin production and increasing gut permeability and inflammation.

**Figure 3.**
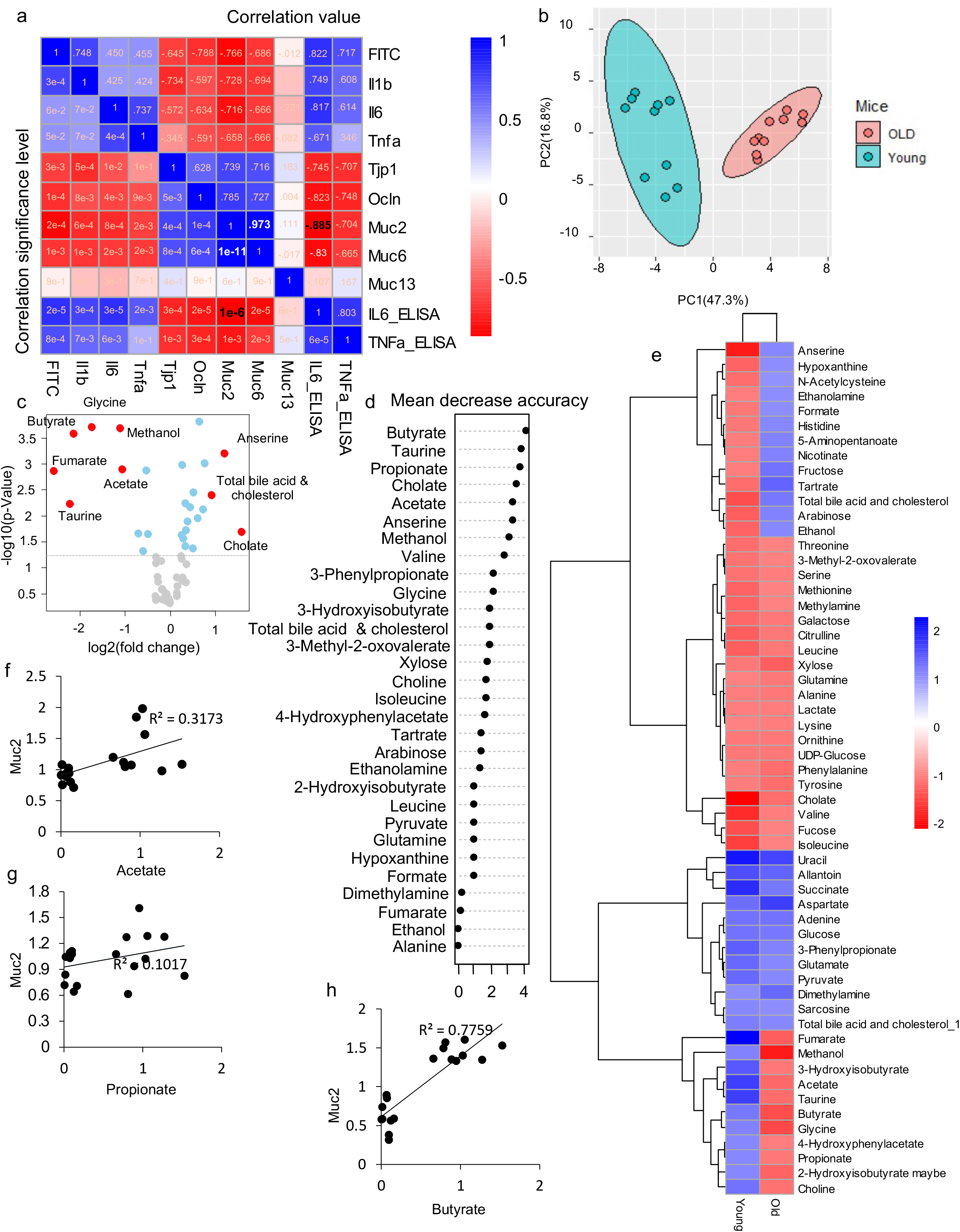
The abundance of short chain fatty acids (SCFAs) like butyrate was significantly reduced in an aged gut as compared to a younger gut, due to reduced mucin barriers. a) Pearson correlation between the mucin, tight junction, leaky gut and inflammatory markers in the gut of aged mice further show Muc2 was dominantly reduced in the aged gut. b,e) Principal component analysis (PCA) (b) and hierarchical differential clustering (e) analyses revealed that an aged gut has significantly different metabolites than a young gut. c,d) Differential abundance analyses of metabolites in volcano plot (c) and random forest analysis show that butyrate was the most significantly reduced metabolite in the aged gut compared to a younger gut. f-h) Further, correlation analyses of Muc2 expression with acetate, propionate and butyrate revealed butyrate was more significantly correlated with Muc2 than acetate and propionate. All the values presented are mean of 5-9 samples in each group.

### 4. Butyrate treatment reduces gut permeability and inflammation by increasing mucus barriers

Feeding butyrate to mice significantly reduced gut permeability and lowered the expression of inflammatory markers (including Il-6, Tnf-α, monocytes chemoattractant protein-1 [Mcp-1] and plasminigen activator inhibitor-1[Pai-1]) (Figure. 4a-b). Further, butyrate treatment significantly increased goblet cell numbers and Muc2 expression in the intestine (ileum) of mice (Figure 4c,d). Butyrate feeding also increased the mucin content in the feces of treated mice compared to non-treated controls (Figure. 4e). These results suggest that butyrate treatment reduced gut permeability and inflammation by increasing goblet cell numbers and mucin production. To determine if butyrate can ameliorate the detrimental effects of aged microbiota (FCM) on mucin barriers, we incubated CMT-93 goblet cells and intestinal organoids with aged gut FCM supplemented with butyrate. Butyrate-incubated CMT-93 cells showed increased mucin production (as suggested by increased PAS staining in Figure 4f) and higher expression of Muc2 and Muc6 compared to controls (Figure 4g). Similarly, the expression of Muc2 and Muc6 was significantly higher and similar to young FCM-treated controls in butyrate-incubated intestinal organoids that were exposed to aged FCM (Figure 4h). These observations showed that butyrate treatment protected the goblet cells from the detrimental effects of the aged microbiota by preserving mucin production. Overall, these results indicate that butyrate reduces aged microbiota-induced disruptions in the mucin barriers mitigating gut permeability and inflammation.

**Figure 4.**
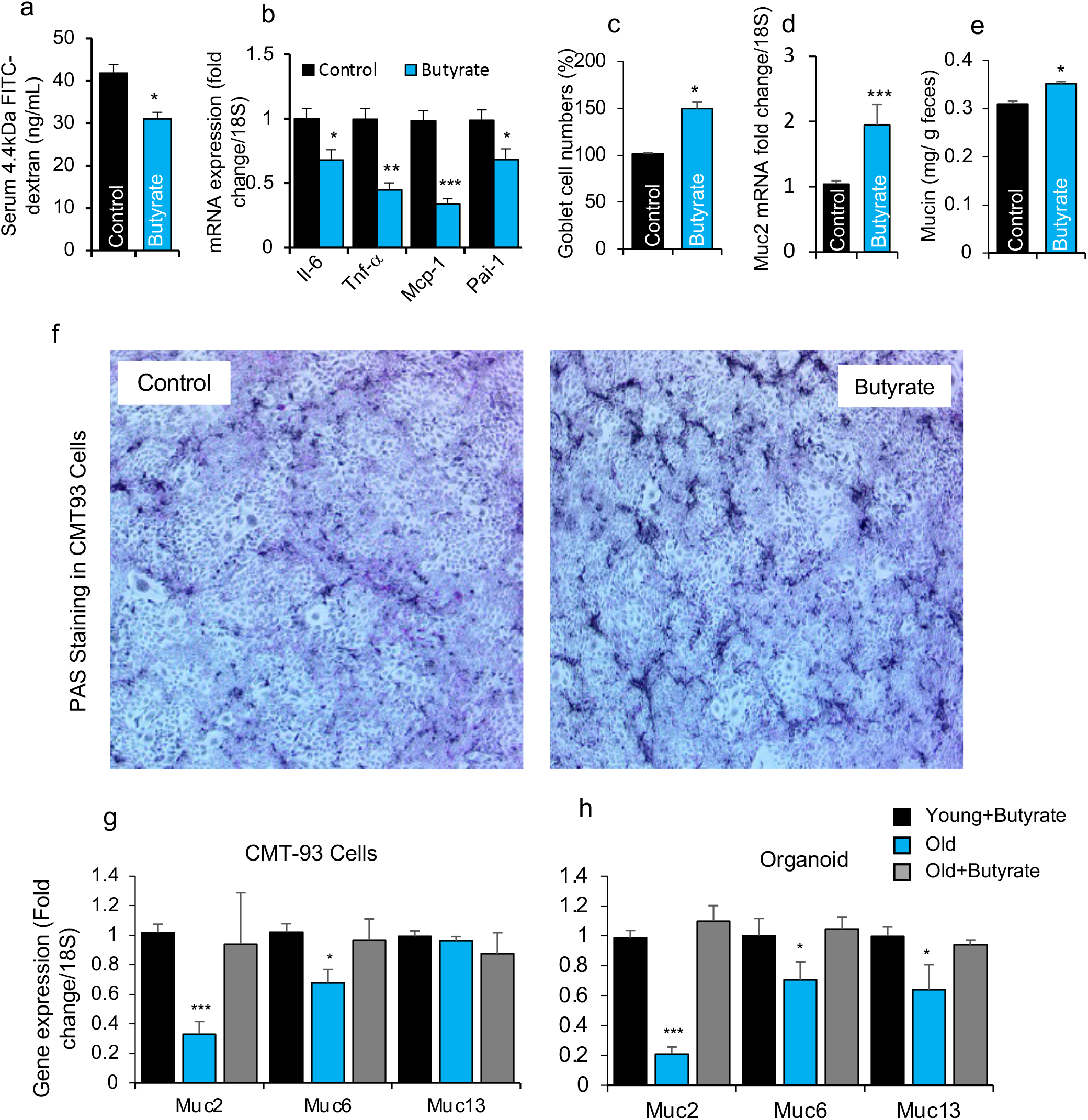
Butyrate treatment decreases leaky gut and inflammation and enhances mucus barriers. a-e) Butyrate feeding in drinking water significantly reduced leakiness (FITC-dextran leakage from gut to blood) (a) and inflammation (IL-6, TNF-α, MCP-1 and PAI-1) (b) and increased goblet cell numbers (c) and expression of Muc2 in the gut along with (d) and mucin in feces of mice fed with a high fat diet as compared to non-butyrate treated controls. e) In addition, butyrate treatment significantly increased mucin production (revealed by Periodic acid– Schiff (PAS) staining) in CMT-93 cells compared to non-treated controls. f-g) Pre-butyrate treatment abrogated detrimental effects of old FCM on the expression of mucin genes in both CMT93 cells (g) and intestinal organoids (h). All the values presented are mean of 5-8 animals, and 3-4 independent replicates in cells and organoid cultures in each group and error bars represent standard error of means. P-values with *p<0.05, **p<0.01, ***p<0.001 are statistically significant. *p<0.05, **p<0.01, ***p<0.001

### 5. Deficiency of SCFAs receptors FFAR2 and FFAR3 increases gut permeability and inflammation via reducing mucin barriers thus resulting in early aging gut phenotype

Although it is known that the SCFAs, like butyrate, act on host cells by activating G-coupled protein receptor signaling (such as FFAR2 [Gpr43] and FFAR3 [Gpr41] (Mishra et al., 2020),,the role of FFAR2/3 in preserving mucin barriers is poorly understood. Expression levels of Ffar2 and Ffar3 were significantly lower in the gut of aged mice compared to younger mice (Figure 5a). Furthermore, expression of Ffar2/3 was significantly decreased in the gut of old FMT recipients as well as in the intestinal organoids, CMT93 and HT29 cells treated with FCM from aged mice compared to young FCM treated controls (Figure 5b-e). These results indicated that the aged microbiota plays a causal role in suppressing the of FFAR2 and FFAR3 expression. Furthermore, the ability of butyrate to induce mucin genes (Muc2, Muc6 and Muc13) was significantly diminished when CMT93 cells and intestinal organoids were treated with Ffar2 inhibitor (CATPB; 1 µM) and Ffar3 inhibitor (Pertussis Toxin; 1 µM) (Figure 5f,g). PAS staining also demonstrated that that butyrate induced mucin production was significantly diminished in CMT93 cells upon treatment with FFAR2 and FFAR3 inhibitors (Figure 5h). Together, these results suggest that stimulation of mucin expression by butyrate is dependent on the activation of FFAR2 and FFAR3 receptors. Further, mice lacking FFAR2 and FFAR3 in their intestine (induced by Villin-Cre) exhibited higher gut permeability and inflammation with reduced expression of mucin and tight junction proteins (Figure 5i-p). These data suggest that deficiency of FFAR2 and FFAR3 results in early age onset gut abnormalities manifested as increased gut permeability and inflammation accompanied with reduced mucin expression. Thus, the aged microbiota suppresses FFAR2/3 signaling, which compromises mucin barrier functions and increases gut permeability. Notably, the suppression of FFAR2/3 signaling attenuates the beneficial effects of butyrate supplementation. In young adult mice, deficiency of FFAR2/3 signaling in the intestine promotes a pathology reminiscent of aged gut, suggesting that suppression of SCFA signaling contributes to the induction of early aging gut phenotype.

**Figure 5.**
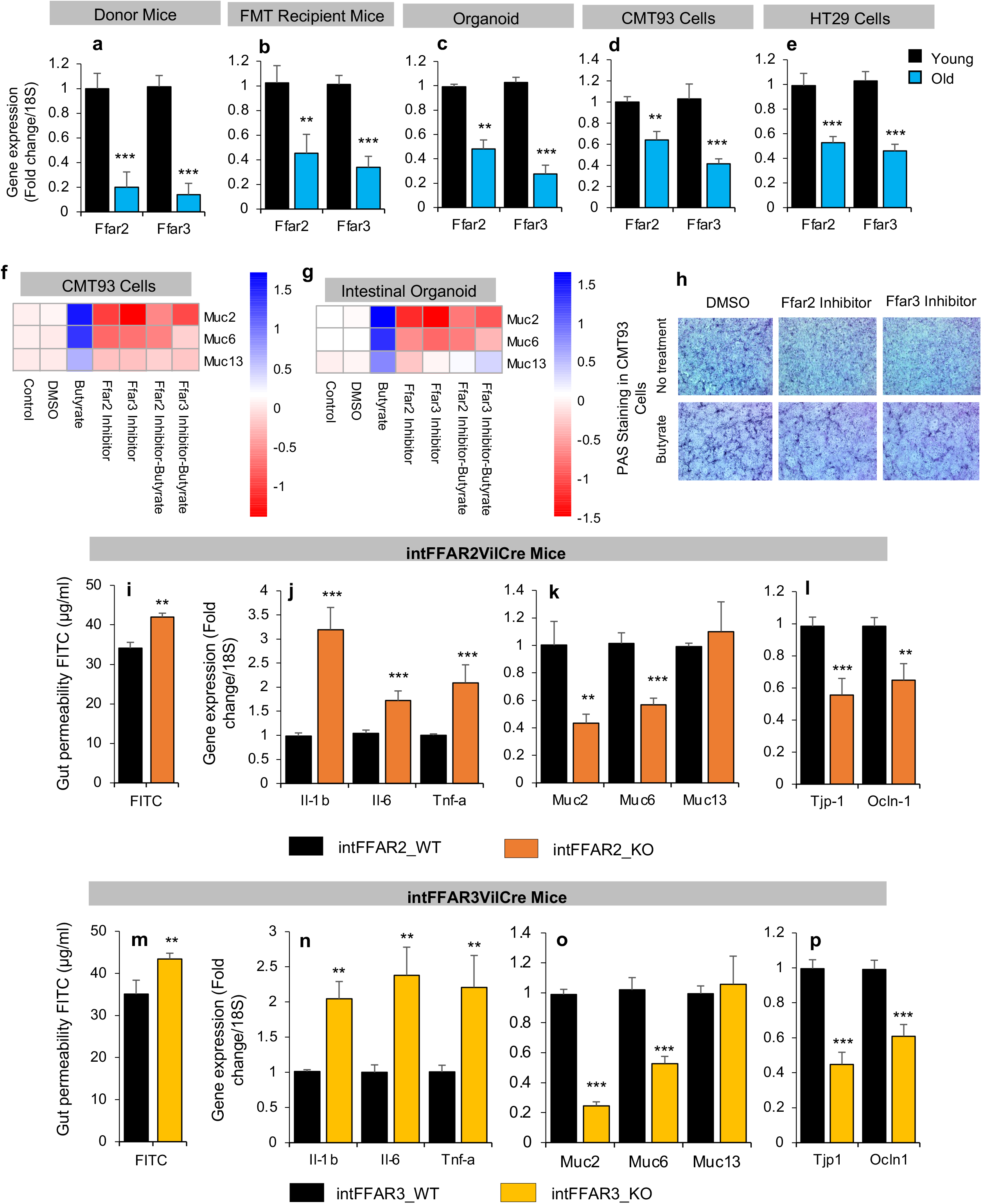
Microbiota reduces FFAR2/3 expression in an aged gut which further depletes butyrate action; Depletion of FFAR2/3 develops early aging in gut. a) The Ffar2 and Ffar3 gene expression was significantly reduced in the gut of old donor mice as compared to young donor mice. b-e) Old FMT and FCM also reduced FFAR2/3 expression in the intestine of the recipient mice gut (b) and treated organoids (c), CMT93 cells (d) and HT29 cells (e) compared to their controls. f-g) Inhibition of FFAR2/3 dampened the effects of butyrate to induce expression of Muc2, Muc6 and Muc13) in CMT93 (f) and intestinal organoids (g), as well as mucin production indicated by PAS staining in CMT93 cells (h). i-p) Intestine specific FFAR2 (intFFAR2 KO) and FFAR3 (intFFAR3 KO) knockout mice show significantly leakier gut (FITC-dextran leakage) (i,m) and expression of inflammatory genes (Il-1β, Il-6 and Tnf-α) (j,n) along with reduced mucin genes (Muc2 and Muc6) (k,o) and tight junction genes (Tjp1 and Ocln) (l,p) in their gut compared to wild type (intFFAR2-WT and intFFAR3-WT) controls. All the values presented are mean of 5-8 animals, and 3-4 independent replicates in cells and organoid cultures in each group and error bars represent standard error of means. P-values with *p<0.05, **p<0.01, ***p<0.001 are statistically significant. *p<0.05, **p<0.01, ***p<0.001

## DISCUSSION

Low grade chronic inflammation is one of the major risk factors of aging-related disorders and, as proposed by the Geroscience hypothesis, stands as one of the key targets to reduce aging related disorders (Kennedy et al., 2014). However, the etiology of inflammation during aging is largely unknown. We and others have reported that aged mice and humans display increased gut permeability (also referred to as leaky gut) that is linked with increased inflammation and poor health (Ferrucci & Fabbri, 2018). Therefore, we reasoned that gut permeability may be an understudied source of inflammation in the aged gut. However, the mechanisms contributing to leakiness in the aged gut remain unknown. Here, we demonstrated that aged gut harbors an abnormal microbiome which causally promotes gut permeability and inflammation by disrupting mucus barriers. We also show that the aged microbiota produces lower amounts of beneficial metabolites like butyrate and suppresses FFAR2/3 signaling resulting in disruption of mucin barriers that promote gut permeability and inflammation.

Inflammation is an immune response that can be induced by microbes. In particular, the non-specific diffusion of microbial antigens from the gut to mucosal and systemic immune systems can stimulate inflammation. Emerging evidence indicates that the gut of aged individuals harbors an abnormal gut microbiome (Tachon et al., 2013). Herein, we demonstrated that the gut microbiota of aged mice was significantly different in comparison to their younger counterparts. However, whether abnormalities in the gut microbiota are merely associated with inflammation or if they play a causal role in the process was unknown. To investigate this, we performed FMT studies in mice and found that abnormal microbiota from the aged gut plays a causal role in gut permeability and inflammation. Specifically, we find that the microbiota from the aged gut directly contributes to the induction of inflammation and gut permeability. We used monolayers of human intestinal cells (HT29) treated by fecal conditioned media (FCM) to confirm these findings and demonstrated that the aged microbiota FCM significantly induced gut permeability. We find that the effects of the aged microbiota on intestinal epithelia are conserved in mice and humans and that these effects can be recapitulated in in-vitro and/or ex-vivo settings.

Gut permeability occurs as a result of structural disruptions in the gut barrier. The mucus layer and tight junction proteins serve as the main components of this barrier (Chelakkot et al., 2018). FMT studies revealed that the aged gut microbiota significantly reduced the expression of mucin and tight junction proteins supporting the notion that the aged microbiota induces gut permeability. Further, unbiased and non-supervised correlational analyses of mice, organoids and human intestinal cells (HT-29) revealed that the *Muc2* gene expression was significantly impacted by the aged microbiota. Mucin is an important biophysical barrier that forms a semipermeable gel like structure on the intestinal epithelial layer to seal and prevent the leakage of gut lumen antigens (Paone & Cani, 2020). However, deterioration of the mucus layer can allow direct contact between the bacterial cells and other antigens in the gut lumen. Such adverse effect on intestinal epithelial cells stimulates the breakdown of tight junction proteins leading to leakage in the gut epithelium.

A common event in mammalian cell interactions with microbial cells (including involving beneficial bacteria) entails an inflammatory response (Belkaid & Hand, 2014). Increased inflammation instigates further breakdown of tight junctions in the intestinal epithelia (Luissint et al., 2016; Zuo et al., 2020). In addition, increased inflammation also reduces production of mucin in the gut (Ahmadi, Razazan, et al., 2020). In aged adults, the mucin layer is thin with this phenotype co-occuring with an abnormal microbiota and increased gut permeability and inflammation. However, if the gut microbiota or inflammation contributes to reducing gut mucin was unclear. Our observations demonstrating that the FMTs of aged microbiota reduced mucin expression in the gut epithelia suggest that microbiota initiatie the disruption of mucin barriers in the gut. However, the reason behind decreased mucin barriers instigated by aged FMTs remained elusive. We here show that old FMTs in mice and treatment of intestinal organoids with old FCMs reduced the number of goblet cells in the intestinal epithelia suggesting that aged microbiota reduces the number of goblet cells. Goblet cells are primarily responsible for mucin production in the gut and further detailed studies are needed to elucidate the mechanism by which the aged microbiota reduces goblet cell numbers in the gut.

To better understand the mechanism of action, we tested the hypothesis that the gut microbiota produces metabolites that impact intestinal epithelial cells (Visconti et al., 2019). To that end, we found that the feces of aged mice exhibited a significantly distinct metabolite signature compared to younger mice, eventhough both groups of mice were fed an identical diet. Furthermore un-biased and non-targeted metabolomic analyses further revealed that the abundance of beneficial metabolites, including SCFAs (such as butyrate), were significantly reduced in the gut of aged mice. Butyrate is a well-studied gut microbiome derived metabolite and considered to be among the most important energy sources for intestinal epithelial cells (Donohoe et al., 2011). Its decrease in the aged gut may result in energy deprivation in intestinal epithelial cells thereby inducing cellular stress. Butyrate levels were positively correlated with mucin expression and butyrate feeding significantly reduced the gut permeability and inflammation via increased mucin production and elevated goblet cell mass. We and others have demonstrated that the feeding of HFD induces gut permeability and inflammation (Ahmadi, Razazan, et al., 2020; Nagpal, Mishra, et al., 2020) by disrupting gut barrier functions (Ahmad et al., 2017). Ahmadi et al also showed that HFD feeding reduces mucin production in aged mice (Ahmadi, Razazan, et al., 2020; Ahmadi, Wang, et al., 2020). However, it is unclear why butyrate levels were decreased in the gut of aged mice. Butyrate levels are often reduced in pathophysiological conditions due to a reduced abundance of butyrate producing bacteria (Jakobsdottir et al., 2013; Lu et al., 2016). A similar mechanism may be involved in aged gut (Lee et al., 2020). On the other hand, it is possible that butyrate uptake is elevated in aged intestinal cells resulting in depletion of butyrate levels.

Although butyrate is one of the most widely studied microbiome metabolites and has important roles in several biological and physiological functions (Liu et al., 2018), its mechanisms of action on mucin biology and gut permeability are poorly understood. Butyrate stimulates G-protein coupled receptors FFAR2 and 3 (also known as Gpr43 and Gpr41) (Mishra et al., 2020). These receptors in turn initiate signaling cascades that target genes involved in gut epithelial cell integrity and inflammation. However, the precise roles of FFAR2 and 3 signaling and its interaction with aged microbiome especially as they pertain to mucin biology and gut permeability are unknown. We showed that expression of FFAR2 and FFAR3 genes were significantly reduced in the aged gut. Further, aged gut microbiota transfer via FMTs or FCM significantly reduced the expression of these genes. Taken together, these findings are consistent with the notion that the aged microbiota suppresses FFAR2/3 expression in the gut. Furthermore, inhibition of FFAR2/3 signaling dampened the beneficial effects of butyrate on mucin expression and production suggesting that FFAR2/3 signaling contributes to butyrate-dependent mucin production. However, SCFAs and butyrate are also known to be taken up and used as an energy source by intestinal cells through infusion into the mitochondrial energy metabolism (NEED ORIGINAL REF OR DELETE THIS) and can also inhibit the activities of histone-deacetylase (NEED ORIGINAL REF OR DELETE THIS). However, the role of these mechanisms in mucin production is not known. Furthermore, intestine specific FFAR2 and FFAR3 knockout mice exhibit significantly low mucin expression along with elevated gut permeability and inflammation at a young age. These findings suggest that compromised FFAR2/3 signaling can recapitulate phenotypes typically seen in the aged gut underscoring the potential importance of this signaling in the early aging phenotype.

Overall, we demonstrate that the aged microbiome promotes gut permeability and inflammation, by disrupting gut barrier functions that are facilitated by a reduction in mucin and goblet cells numbers. The aged gut microbiota reduces butyrate synthesis and FFAR2/3 signaling which, in turn, decreases mucin formation thus weakening gut barrier functions to promote gut permeability and inflammation. Importantly, butyrate administration reverses the phenotype via activation of FFAR2/3 signaling, suggesting that the butyrate supplementation and/or FFAR2/3 agonism represent an effective approach for reversing microbiome abnormalities, gut permeability and inflammation typically seen in aging.

## Material and Methods

### Chemical, reagents, and instruments

The specific information about the chemicals, reagents and instruments including catalog numbers and vendors is provided in Supplementary Table S5.

### Mice studies

Twelve-week (young donor, n=10) and 78 week (aged donor, n=10)C57BL/6J (B6) male mice were purchased from Jackson Laboratory (Bar Harbor, ME, USA) and were acclimatized for 2 weeks in the animal vivarium under a 12h light/-dark cycle. After acclimatization, fresh fecal samples were collected, snap frozen and stored in the −80°C freezer, and were used for 16S metagenomics, metabolomics, and FMT and FCM studies, as described below. Eight week old B6 mice received FMTs from young and old donors, after 7 days of FMT, gut permeability assay was performed and mice were euthanized, and tissues were collected for further analysis. Because high fat diet (HFD) rapidly progresses leaky gut in aging models, some B6 mice were given butyrate (2.5% solution) in their normal drinking water (n=6) while controls were given normal drinking water (n=6), while both groups were fed with high fat diet (60% kcal fat). Homozygous Ffar2^flox/flox (fl/fl)^ and Ffar3^fl/fl^ mice were breed with Vil^Cre+^ heterozygotes mice to develop the intestinal tissue specific -Ffar2 knockout mice (intFFAR2 KO [Ffar2^fl/fl^Vil^Cre+^]), -Ffar3 knock-out mice (intFFAR3 KO [FFAR3^fl/fl^/Vil^Cre+^]) and compared with their corresponding wild type counterparts like intFFAR2 WT (FFAR2^fl/fl^-Vil^Cre-^) and Ffar3 (Ffar3^fl/fl^Vil^Cre-^) at the 10 weeks of age for gut permeability, inflammation and mucus barriers while fed with normal chow. All the animal experiments and procedures were approved by the IACUC of the Wake Forest School of Medicine and University of South Florida.

### Fecal microbiome transplantation (FMT)

We have used gut cleansed mice instead of germ-free mice in our studies, because germ free mice have several abnormalities in the gut which may confound the results of our studies. To perform gutcleansing in the recipient mice, we gave them an antibiotic cocktail (consisting of 1 g/l Ampicillin, 1 g/l Metronidazole, 1 g/l Neomycin, and 0.5 g/l Vancomycin) for 4 days in the drinking water. On the 4^th^ day, all the mice were fasted for 4 hours and 4 doses of polyethylene glycol (PEG) were given at 20 minutes interval using oral gavage. PEG physically cleanses the gut and reduces upto the 95% of microbes. After cleansing, the fecal slurry (200 uL) was prepared from the donor groups/for the recipients groups. We have transplanted 9 mice with the young microbiota and 9 mice with old microbiota (Ahmadi, Razazan, et al., 2020; Wang et al., 2020; Wrzosek et al., 2018). We continued giving one dose of fecal slurry every day for an additional 4 days to stabilize the microbiome and maintain it closer to the donor microbiome. The fecal slurry was prepared using snap frozen fecal samples (500 mg) from donor mice, which were dissolved in 5 ml of reduced PBS (phosphate buffer supplemented with 0.1% Resazurin (w/v) and 0.05% L-cysteine-HCl in anaerobic chamber (Kang et al., 2017).

### Gut permeability assay

Mice were pre-fasted for 4 hours and administered with 4-kD FITC (Fluorescein isothiocyanate)-dextran at a concentration of 1g/kg body weight through oral gavage. After, an additional 4 hours of fasting, blood serum was collected from the tail vein to measure the appearance of FITC fluorescence at 485 nm excitation and at 530 nm emission wavelength using a fluorescence 96-well plate. The FITC concentration was determined using a standard FITC curve as previously described in our earlier publications (Ahmadi et al., 2019; Ahmadi, Razazan, et al., 2020; Ahmadi, Wang, et al., 2020; Nagpal, Newman, et al., 2018; Wang et al., 2020).

### Markers of systemic leaky gut and inflammation assays

The concentrations of LBP and sCD14 were measured in mice serum to determine the systemic leaky gut; while the concentration of IL-6 and TNF-α were measured in mice serum to determine systemic inflammation using the ELISA kits.

### Fecal mucin content

Fecal mucin content was quantified using an ELISA kit following the manufacturer’s protocol. Mucin content was measured in triplicate from each mice feces.

### Histochemistry

The intestinal tissues were collected, washed with PBS, and fixed in 10% formalin. Tissues were sliced into 8-µm thick sections, which were stained with hematoxylin and eosin (H&E). Goblet cells were counted as unstained vacuoles in the H&E immunohistochemistry slides. Mucus staining in the goblet cell line- CMT93 cells was performed in a 12-well cell culture plate by using the Alcian Blue/PAS kit following the manufacturer instructions. All the images were captured at 20X magnification using AmScope microscope.

### Gut microbiota analyses

Our well established 16S rRNA sequencing with the MiSeq Illumina platform and bioinformatics were used to analyze the gut microbiota composition (Ahmadi et al., 2019; Nagpal, Mishra, et al., 2020; Nagpal et al., 2019; Nagpal, Neth, et al., 2020; Nagpal, Shively, et al., 2018; Nagpal, Wang, et al., 2018). In brief, genomic DNA was extracted from ∼100 mg of mice feces using the Qiagen QiaAmp PowerFecal Pro kit. Primers 515 F (barcoded) and 806 R were used to amplify the V4 region of bacterial 16S rDNA (Caporaso et al., 2010). After being purified and quantified with AMPure® magnetic purification beads (Agencourt) and Qubit-3 fluorimeter, respectively. Equal amounts (8 pM) of the amplicons were used for sequencing on an Illumina MiSeq sequencer (using Miseq reagent kit v3). The sequences were de-multiplexed, quality filtered, clustered, and analyzed using the Quantitative Insights into Microbial Ecology (QIIME) and R-based analytical tools (Navas-Molina et al., 2013).

### Metabolomics analysis

Fecal samples were extracted using a previously described method (Gratton et al., 2016) with slight modifications. The extracted water samples were mixed with phosphate buffer and the final solutions contain 10% of D_2_O with 0.1 M phosphate buffer (pH = 7.4) and 0.5 mM trimethylsilylpropanoic acid (TSP). NMR experiments were conducted on a Bruker Ascend 400 MHz high-resolution NMR using a 1D first increment of a NOESY (noesygppr1d) with water suppression and a 4-s recycle delay. All NMR spectra were phased and referenced to TSP in TopSpin 4.06 (Bruker BioSpin, Ettlingen, Germany). The NMR peaks were extracted using Amix 4.0 (Bruker Biospin, Ettlingen, Germany) with a previously reported automatic bucketing method to minimize peak splitting. The metabolite identification was carried out using Chenomx 8.6 (Chenomx Inc, Edmonton, Alberta Canada). Total intensity normalization was applied before further data analysis.

### Preparation of Fecal Condition Media (FCM)

The snap frozen feces were ground in liquid nitrogen using a mortar and pestle. The fine powdered feces were suspended in cold Dulbecco’s modified eagle’s medium (DMEM)media at a concentration of 100 mg fecal powdered content in 100 ml of media. The suspended fecal media was shaken at a speed of 200 rpm for 1 hour at 4°C to mix properly. The suspended media was filtered twice through a 0.45 µm pore nylon membrane sterile filter and twice through a 0.22 µm nylon membrane sterile syringe filter in sterile condition. This prepared FCM with 1:40 dilution was used to challenge the organoid and cells at a concentration of 2.5 mg/ml.

### Intestinal organoid development

The intestinal organoids were prepared as described earlier by us (Nagpal, Newman, et al., 2018). In brief, the entire small intestine of 6-8 week old B6 mice was washed with pre-cooled Dulbecco’s phosphate buffered saline (DPBS), and fragmented into the 2 mm small pieces after being cut opened lengthwise. These tissue fragments were washed gently with pre-cooled PBS by pipetting up and down for next 15-20 times until the supernatant become clear.Then, the fragments were suspended in 25 ml of pre-warmed trypsin and shaken for 15 minutes at room temperature. Trypsin was removed by adding four times of fresh 10 ml pre-cold PBS with 0.1% bovine serum albumin (BSA). Digested fragments were filtered through a 70 μm cell strainer and washed by centrifuging at 290x g for 5 min at 4°C. This process was repeated two more times. The intestinal crypt-containing pellets were resuspended in 10 mL of cold DMEM: Nutrient Mixture F-12 (DMEM/F12) and centrifuged at 500x g for 10 minutes at 4 °C before resuspension into 150 μL IntestiCult Organoid Growth Medium containing with 50 μg/mL gentamicin. The Matrigel Matrix was added to the suspension and 50 μL of the mixture was pipetted slowly at the center of a prewarmed 24-well culture plate to form a dome. The plate was immediately incubated at 37°C and 5% CO_2_ for 30 minutes to allow the Matrigel to set, and 500 μL of IntestiCult Organoid Growth Medium was added into each well. To maintain the cultures, the IntestiCult Organoid Growth Medium was changed three times per week. After 48 hours, the developed organoids were challenged with Ffar2 inhibitor (1 µM CATBP) and Ffar3 inhibitor (1 µM Pertussis Toxin) and on day 10, the organoid were challenged with young and old FCM (2.5 mg/ml) and butyrate (6 µM). The organoids were harvested on 12^th^ day for gene expression analyses. Each experiment was conducted 3 times with 3 replica each.

### Cell Culture

Human HT-29 cells and mouse CMT-93 were purchased from American Type Culture Collection (ATCC). The HT-29 cells (passages 10-15) and CMT93 cells (passages 6-14) were maintained in 4.5 g/L D-Glucose with L-Glutamine DMEM supplemented with 10% fetal bovine serum (FBS) and 100 U/mL penicillin, 100 U/mL streptomycin. The CMT93 cells were grown for 48 hours then treated with young and old FCM (2.5 mg/ml), butyrate (6 µM), Ffar2 inhibitor (1 µM CATBP) and Ffar3 inhibitor (1 µM Pertussis Toxin) for 12 hours. Fully differentiated 21-days HT-29 cells were used for young and old FCM treatment (2.5 mg/ml) for a time duration of 8 hours to measure the TEER and FITC diffusion assay. The cells were washed properly with DPBS before harvesting.

### Measurements of transepithelial electrical resistance (TEER) in HT-29 Cells

Caco-2 cells were seeded on apical chamber made of polyester membrane filters with 0.4 μm pore size of 12-well transwell plates at a density of 3 × 10^5^/well. The culture media from both the apical and basolateral compartments was changed every two days. The cells were allowed to fully differentiate for the next 21 days. On the 21^st^ day, the fully differentiated cells were challenged with FCM for 8 hours with continuous measuring of TEER values using an EVOM^2^ Epithelial Voltohmmeter according to the manufacturer’s instruction. The blank inert resistance value (the insert with only culture media) was subtracted from the measured resistance value of each sample and the final resistance in (reported in ohm×cm^2^) was calculated by multiplying the sample resistance by the area of the membrane(Ahmadi, Wang, et al., 2020).

### FITC-dextran permeability assay in HT-29 cells

The HT-29 cell transwell plates were prepared using the same method as in the TEER experiments. On the 21^st^ day, the HT-29 cells monolayer was treated with FCM and 4-kD-FITC-dextran solution (1 mg/ml) was added on the apical (upper) side of the monolayers.The 4-kD-FITC level of the basolateral side was determined using a fluorescent 96-well [late at a 485 nm excitation and at 525 nm emission wavelength using fluorescence 96-well plate reader. Values were normalized using a standard FITC curve as described in our paper(Wang et al., 2020).The assay was conducted three times in triplicate.

### Gene expression using real-time PCR

Total RNA was extracted using RNeasy Mini Kit following the manufacturing protocol from frozen tissues, intestinal organoids, and cells. cDNA was synthesized using High-Capacity cDNA reverse transcription kit. The gene expression was quantified using ABI 7500 real time PCR machine using powerUp SYBR Green master mix using specific primers (listed in Supplementary Table S3 and S4). Relative gene expression was analyzed using ^ΔΔ^CT method by normalizing with 18S as an internal housekeeping control (Ahmadi, Razazan, et al., 2020; Ahmadi, Wang, et al., 2020; Yadav et al., 2017; Yadav et al., 2013).

### Statistical Analyses

Different datasets were analyzed using the student’s t-test. All the statistically analyzed figures are presented in the form of mean ± standard error of mean (SEM). Alpha-diversity indices and the bacterial abundance between the old versus young was compared using the unpaired two-tailed Student’s t-test. Differences in beta-diversity were tested by permutational multivariate analysis of variance (PERMANOVA), a permutation-based multivariate analysis of variance to a matrix of pairwise distance to partition the inter-group and intra-group distance. Random forest analysis (RFA) and principal component analysis (PCA) were analysed in R programming (version 3.6.0 https://www.r-project.org/) using packages “randomForest”, “ggplot2”, “caret”, “psych”, “ggbiplots”, “nnet” and “devtools”. Hierarchical clustering and heatmaps of top 50 genera based on abundance was constructed by using the pheatmap and “ggplots” packages of R v6.0. LEfSE (Linear discriminatory analysis [LDA] Effect Size) was used to identify unique bacterial taxa that drive differences in old vs young mice fecal sample from Galaxy server (https://huttenhower.sph.harvard.edu/galaxy/) (Segata et al., 2011). The alpha parameter significance threshold for the Kruskal–Wallis as well as the Wilcoxon signed-rank test, which were implemented among classes was set to 0.01, the logarithmic LDA score cut-off was set to 3, and the strategy for multi-class analysis was set to “all-against-all”. Welch’s t-test was applied for statistical significance analysis for metabolites using Amix 3.9 (Bruker Biospin) and a false discovery rate (FDR) were applied to control the family wised error. Hierarchical clustering and heat-maps were constructed based on the average linkage on Euclidean distance and depict the patterns of abundance and log values of metabolites and the correlation of these metabolites with gut permeability, inflammatory and tight junction marker were constructedin R v6.0 using the pheatmap and “ggplots” packages. The Pearson’s correlation between gut permeability, inflammatory and tight junction markers were calculated between respective groups in IBM SPSS (statistical package for social science) and clustering correlation heatmaps were constructed using the pheatmap package, R v6.0. P<0.05 was considered statistically significant for all the analyses unless otherwise specified.

## Acknowledgement

All the authors are grateful for all the fellow staff and laboratory members of Yadav’s lab for their cooperation and help during the study.

## Funding

We are thankful for the support provided by National Institutes of Health (NIH) grants R56AG064075, R56AG069676, R21AG072379, and the Department of Defense funding W81XWH-19-1-0236 (HY), as well funds and services provided from the USF Center for Microbiome Research and Center for Excellence in Aging and Brain Repair.

## Declaration of conflict of interest

All authors declare no conflict of interest for this work.

## SUPPLEMENTARY FIGURES AND FIGURE LEGENDS

**Supplementary Figure S1.**
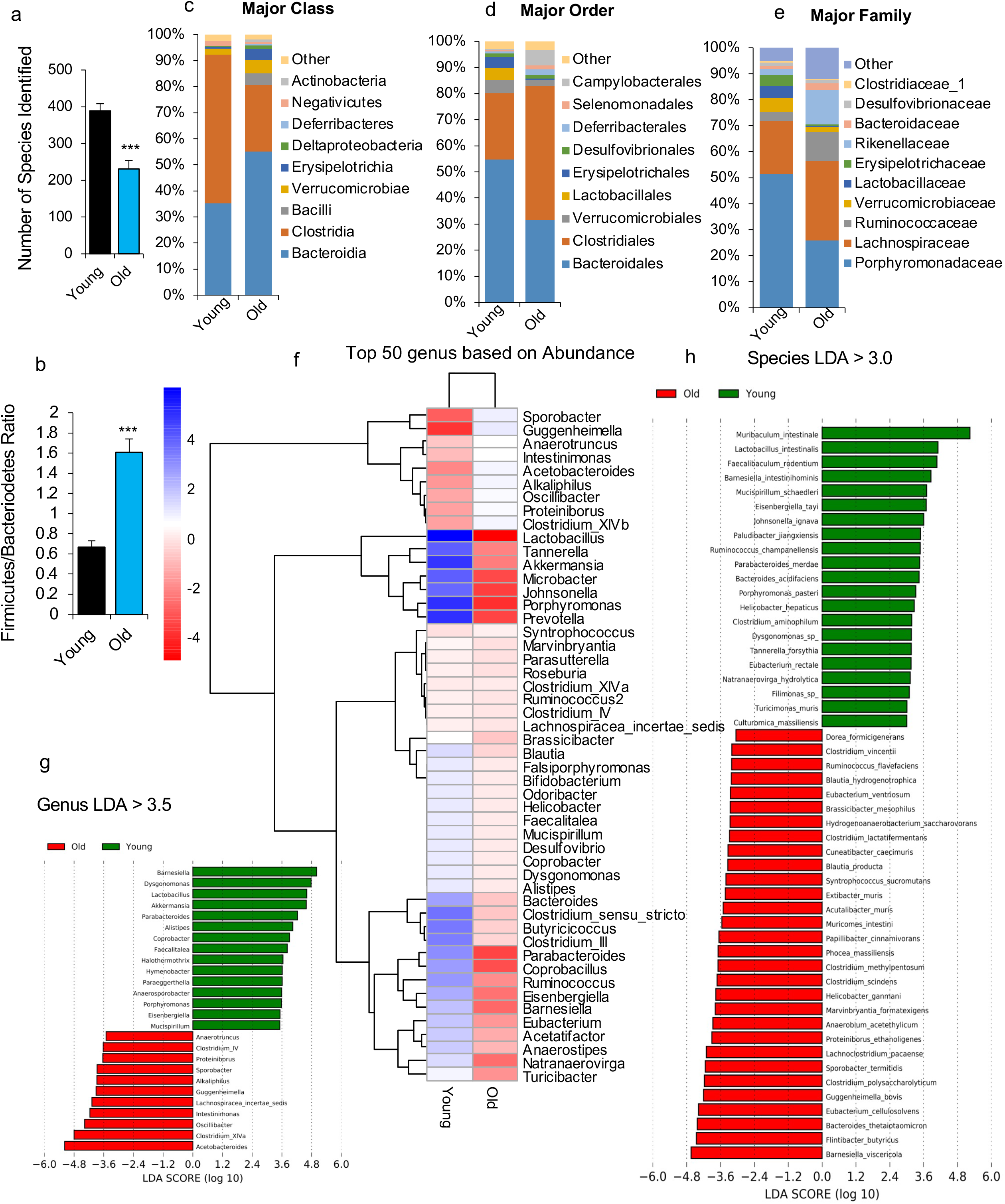
Microbiota in aged donor mice is significantly distinct from young donor mice. a) Alpha diversity indices show the number of observed is significantly lower in the feces of aged mice compared to younger mice. b) The ratio of Firmicutes/Bacteroidetes was significantly higher in aged mice feces compared to younger counterparts. c-e) The abundance of major class (c), order (d) and family (e) of bacteria that are distinct in aged versus younger gut. f) Hierarchical heatmap of top 50 genus based on abundance in old vs young mice feces. g-h) Linear discriminant analysis (LDA) effect size LEfSe of old microbiome at genera and species level as compared to younger microbiome. All the values presented are mean of 5-9 animals in each group and error bars represent standard error of means. P-values with ***p<0.001 are statistically significant.

**Supplementary Figure S2.**
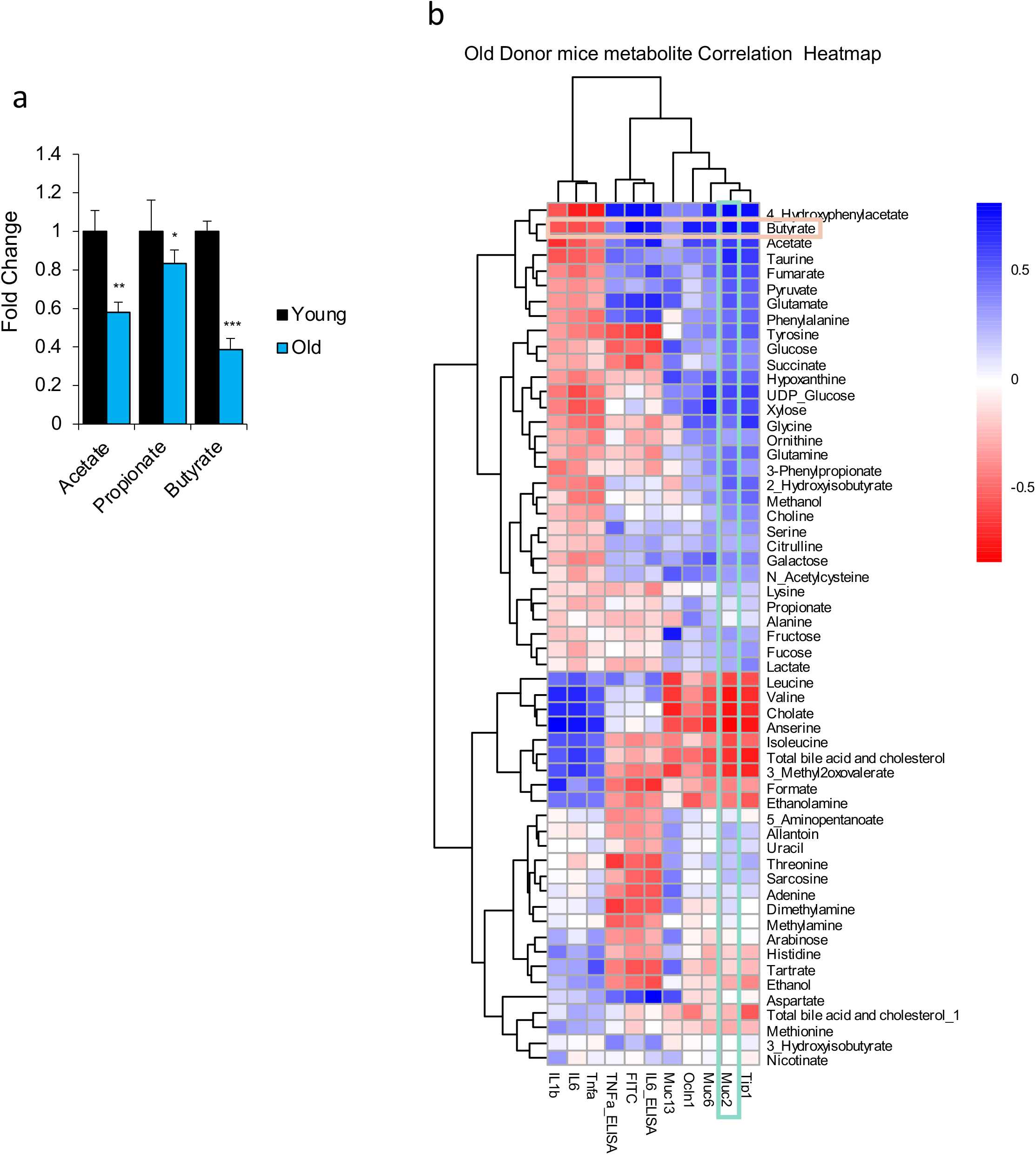
The abundance of metabolites was significant different in aged versus younger gut and are distinctly correlated with mucin, tight junctions and leaky and inflammatory markers. a) The abundance of SCFAs (acetate, propionate and butyrate) was significantly lower in the feces of aged mice compared to younger mice. b) Heatmap of Pearson correlation values among microbiome metabolites and mucin, tight junction protein, leaky gut and inflammatory markers revealed significantly unique clustering, with the highest correction in butyrate and Muc2 gene expression.

## Notes

### Competing Interest Statement

The authors have declared no competing interest.

